# Multiplex chromatin interaction analysis by signal processing and statistical algorithms

**DOI:** 10.1101/665232

**Authors:** Minji Kim, Meizhen Zheng, Simon Zhongyuan Tian, Daniel Capurso, Byoungkoo Lee, Jeffrey H. Chuang, Yijun Ruan

## Abstract

The single-molecule multiplex chromatin interaction data generated by emerging non-ligation-based 3D genome mapping technologies provide novel insights into high dimensional chromatin organization, yet introduce new computational challenges. We developed MIA-Sig (https://github.com/TheJacksonLaboratory/mia-sig.git), an algorithmic framework to de-noise the data, assess the statistical significance of chromatin complexes, and identify topological domains and inter-domain contacts. On chromatin immunoprecipitation (ChIP)-enriched data, MIA-Sig can clearly distinguish the protein-associated interactions from the non-specific topological domains.

## Main text

Previous 3D genome-mapping efforts have suggested complex chromosomal folding structures. In particular, methods based on high-throughput sequencing capture bulk chromatin contacts (Hi-C (Lieberman-Aiden et al., 2009)) or enrich for chromatin contacts involving a specific protein (ChIA-PET (Fullwood et al., 2009)). Both of these methods rely on proximity ligation, and therefore can only reveal population averages of pairwise contacts. Thus, they lacked the ability to simultaneously capture multiple loci involved in a chromatin complex in an individual cell.

To overcome these challenges, novel experimental methods have recently been developed to capture multiplex chromatin contacts with single-molecule resolution. For instance, GAM (Beagrie et al., 2017) identifies multi-way interactions by capturing multiple DNA elements co-existing in a given nuclear slice, SPRITE (Quinodoz et al., 2018) barcodes individual chromatin complexes via a split-pool strategy, and ChIA-Drop (Zheng et al., 2019) partitions each complex into a microfluidic droplets for barcoding and amplification. Collectively, these emerging 3D genome-mapping technologies are advancing the frontier of the nuclear architecture field. However, as with other genomic approaches prone to the background noise, the noisy and high-dimensional nature of the multiplex data poses unique computational challenges that cannot be readily addressed with existing tools that are tailored for pairwise interactions data. Thus, we developed MIA-Sig (<u>M</u>ultiplex <u>I</u>nteractions <u>A</u>nalysis by <u>S</u>ignal processing algorithms) with a set of Python modules tailored for ChIA-Drop and related datatypes.

A central analytic challenge is to distinguish the true biological chromatin complexes from the experimental noise. A possible source of noise is an event that two or more chromatin complexes are potentially encapsulated in the same microfluidics droplet and then assigned the same barcode, yielding a multiplet (**Figure 1a**). The problem also prevails in microfluidics-based single-cell RNA-seq data, which is then resolved computationally via dimensionality-reduction and clustering (Wolock et al., 2019). However, methods developed for single-cell transcriptomics data are not apt for multiplex chromatin interactions data since: 1) the signal for chromatin interactions is point data (fragment is captured or not captured) rather than continuously valued data (gene expression level), and 2) multiplex chromatin interaction data are inherently sparser than the single-cell transcriptomics data, due to the lack of cell barcodes.

**Figure 1:**
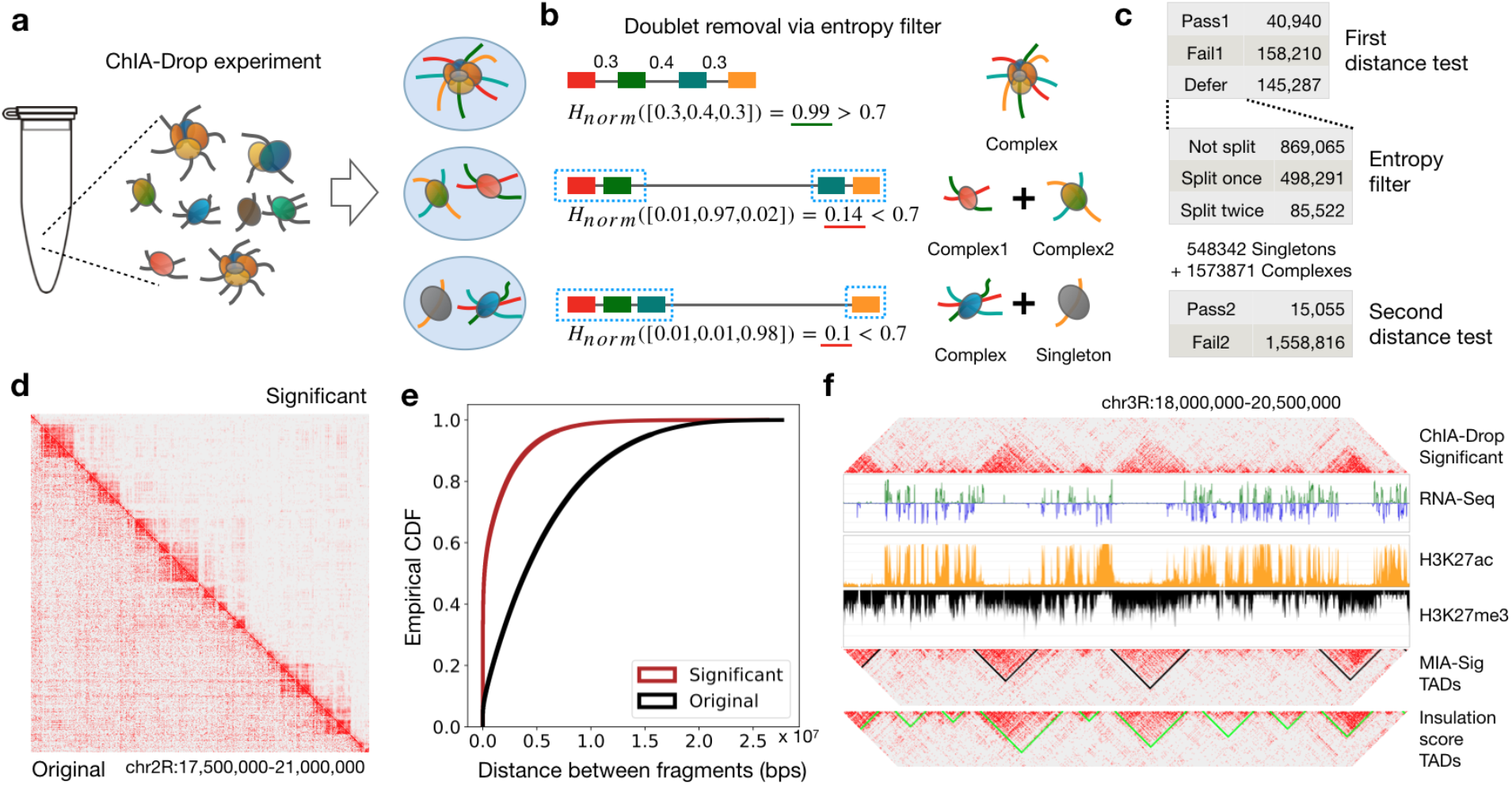
Performance of MIA-Sig on ChIA-Drop data. (a) ChIA-Drop experiments are designed to encapsulate each chromatin complex in a droplet, but the encapsulation is a random process and sometimes results in more than one complex in a droplet (multiplets). (b) MIA-Sig aims to detect multiplets by computing the normalized Shannon entropy *H_norm_* (Methods). It separates a complex at the largest distance if *H_norm_* is smaller than a threshold, which is 0.7 in this example. (c) Summary statistics of the distance test indicate that entropy filter resolves around 500,000 doublets and 85,000 triplets, from which 15,055 complexes pass the second distance test. (d) 2D heatmap comparison of original (bottom triangle) and significant (upper triangle) complexes demonstrates that MIA-Sig removes off-diagonal noise. (e) Empirical cumulative distribution function for the neighboring distances of original and significant complexes (two-sided Kolmogorov-Smirnov test statistic= 0.47, p-value < 2.2 × 10^-16^). (f) Comparison of TADs called from significant putative complexes (MIA-Sig) and from enumerating all pairs of fragments (Insulation score). MIA-Sig more specifically separates active regions (high H3K27ac and low H3K27me3) rather than assigning them to TADs.

Therefore, we devised a distance test with an entropy filter based on the biological knowledge that most meaningful chromatin interactions occur in a certain distance range, while those outside the range are likely noise (Lajoie et al., 2015). By converting the distances between fragments into a probability vector, we compute the normalized Shannon entropy (Shannon, 1948), ranging from 0 to 1. If a droplet contains a single complex, the fragments are presumably close and equally spaced, resulting in high entropy close to 1. In the case of a doublet, two independent complexes would be separated by a single large distance, resulting in low entropy close to 0, which can then be separated into two singlets (**Figure 1b**). To identify significant chromatin complexes, a resampling-based distance test is applied before and after the entropy filter (**Figure 1c**; **Methods**). As a result, we retained 55,995 statistically significant complexes in the Drosophila S2 ChIA-Drop data out of 3,075,926 putative complexes (**Supplementary Figure 1**). Filtering to retain significant complexes preserves the TADs along the diagonal of the 2D heat maps, while reducing the off-diagonal noise (**Figure 1d**; visualization through Juicebox (Durand et al., 2016)). A shift in distance distributions supports that meaningful interactions are captured within 10 kb and 1 Mb (**Figure 1e; Supplementary Figure 2**). Of the significant chromatin complexes, 15,055 (27%) were from the entropy filtering step that resolved doublets and triplets (**Supplementary Figure 3**).

From the significant complexes, it is desirable to automatically call TADs for downstream analyses. Many TAD calling algorithms exist for Hi-C data (Zufferey et al., 2018), yet all are based on pairwise contacts. To retain multiplexity information, we developed an algorithm to call TADs directly from ChIA-Drop data (**Methods**). The idea is to convert complexes into 1D signal track, then apply wavelet transformation (Mallat, 1989) to smooth the signal while retaining clear change points (**Supplementary Figure 4**). MIA-Sig called 335 TADs with wider range of sizes than 513 TADs called by pairwise ‘Insulation Score’ (‘InS’) approach (**Supplementary Figure 5**). Compared to InS TADs, the MIA-Sig TADs are less likely to overlap active regions characterized by high H3K27ac and low H3K27me3 (**Figure 1f**), which are known to be the gaps between TADs in *Drosophila* (Rowley et al., 2017). This pattern is observed genome-wide: MIA-Sig TADs have higher inactive mark (H3K27me3) in than InS TADs, and MIA-Sig gaps have higher active mark (H3K27ac) than InS gaps (**Supplementary Figure 6**). Most interactions occur within a single TAD, but 23% of significant complexes also cross 2 or more TADs, consistent with previous findings (Paulsen et al., 2019). Thus, we identify frequent interactions involving multiple TADs by counting the occurrences and performing a binomial test (**Methods**). A set of TADs with frequent contacts are ultimately assigned low p-values (**Supplementary Figure 7**), which can guide the researchers to perform validation experiments.

Similar to ChIA-PET, ChIA-Drop can also enrich chromatin complexes involving a specific protein, such as RNAPII or CTCF. We implemented an enrichment test to estimate the significance of binding intensity of observed chromatin complexes (**Figure 2a; Methods**) compared to an empirical null distribution (**Supplementary Figure 8**). As a result, we retain significant complexes with their fragments in highly enriched domains characterized by high RNA-seq expression and H3K27ac signal with abundant RNAPII ChIA-PET loops (**Figure 2b**). Genome-wide patterns confirm that significant complexes are biased towards active regions, whereas insignificant complexes are not (**Supplementary Figure 9**). Moreover, significant complexes have higher median H3K27ac signals and lower median H3K27me3 signals than insignificant complexes (**Figure 2c-d**). A detailed view around a few genes shows that significant complexes are more likely to retain promoter-centric interactions than insignificant complexes (**Figure 2d**; visualization through ChIA-View (Tian et al., 2019)). This pattern is prevalent genome-wide, with 69% of significant complexes containing at least 1 promoter compared to only 30% of insignificant complexes (**Figure 2f**). Notably, significant complexes are most likely to capture one active promoter and one or more non-promoters—possibly enhancers—while insignificant complexes are prone to detect interactions among non-promoters (**Supplementary Figure 10**). Among the promoter-involving fragments, those in significant complexes have higher median gene expression than those in insignificant ones.

**Figure 2:**
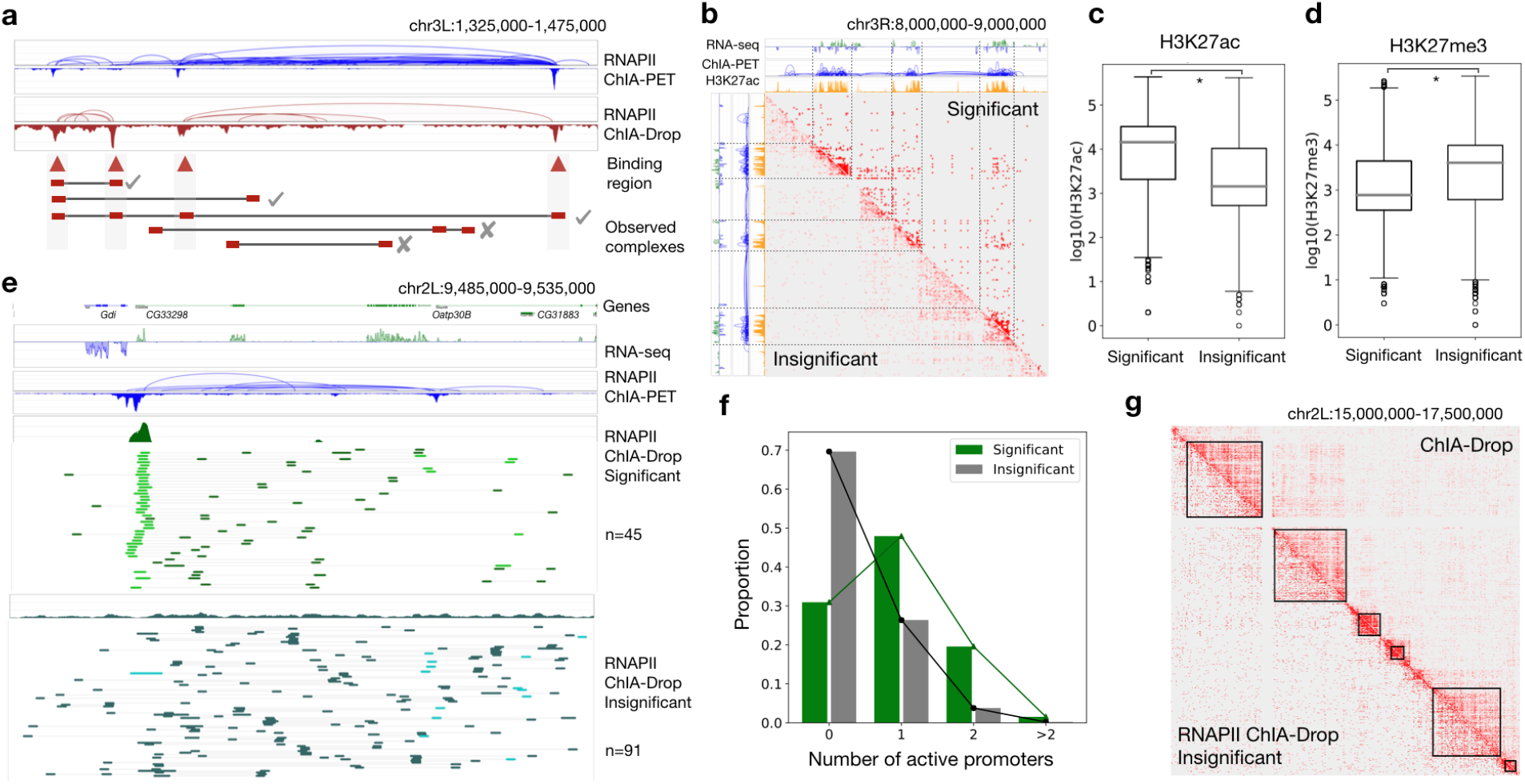
Enrichment test on RNAPII ChIA-Drop data. (a) MIA-Sig performs an enrichment test on RNAPII-enriched ChIA-Drop data by retaining complexes with fragments in strong binding regions, which also correspond to RNAPII ChIA-PET peaks. (b) The significant complexes are pronounced in regions with high level of transcription, abundant loops, and active histone mark; insignificant complexes tend to be in inactive regions. (c) Log of H3K27ac signal for fragments in significant and insignificant complexes (one-sided Mann-Whitney U test, p-value < 2.2 × 10^-16^). (d) Log of H3K27me3 signal for fragments in significant and insignificant complexes (one-sided Mann-Whitney U test, p-value < 2.2 × 10^-16^). (e) Fragment coverage profile of significant complexes is similar to that of RNAPII ChIA-PET, with 45 promoter-centric multiplex interactions (green: non-promoters, light green: promoters). By contrast, insignificant complexes do not show any strong binding peaks in coverage, and 91 multiplex interactions are non-specific (turquoise: non-promoters, light turquoise: promoters). (f) Genome-wide, significant complexes have higher proportion of active promoter fragments than insignificant complexes do (two-sided K-S test statistic= 0.39, p-value < 2.2 × 10^-16^). (g) The insignificant RNAPII ChIA-Drop complexes from the enrichment test are comparable to the significant ChIA-Drop complexes from the distance test. TADs (black lines) are called by MIA-Sig on the latter complexes.

As with many experimental protocols, the chromatin immunoprecipitation step is not 100% efficient and typically yields a 20-40% efficiency rate (Tang et al., 2015). Thus, we take advantage of the fact that enriched ChIA-Drop data sets also contain some background signal for chromatin complexes that did not specifically involve the protein of interest, similar to nonenriched ChIA-Drop data. Through the MIA-Sig enrichment test on RNAPII ChIA-Drop data, we can extract the non-enriched complexes from the insignificant complexes, which approximately emulate the ChIA-Drop data (**Figure 2g**).

Given that the frontier of the nuclear architecture field is now single-cell and singlemolecule 3D genome mapping, it is imperative to develop algorithms to analyze data from these novel experimental protocols. We have presented an approach to the imminent problem of extracting statistically significant complexes from noisy signals, calling TADs, and identifying frequent inter-TAD contacts (**Figure 3**). In addition, we offer a practical strategy to extract nonenriched ChIA-Drop from RNAPII ChIA-Drop. As a publicly available software package, MIA-Sig provides a valuable algorithmic framework for multiplex chromatin interaction data to be utilized by the broader scientific community. MIA-Sig nonetheless has its own drawbacks. One key assumption in the distance test is that a fragment far from the other fragments is likely a droplet contamination resulting in a doublet, a behavior yet to be confirmed experimentally and statistically. In performing the enrichment test for RNAPII ChIA-Drop data, we do not use a background distribution model and instead draw an empirical null distribution via random sampling. A disadvantage of this approach is the computational cost, which can be demanding for large human datasets.

**Figure 3:**
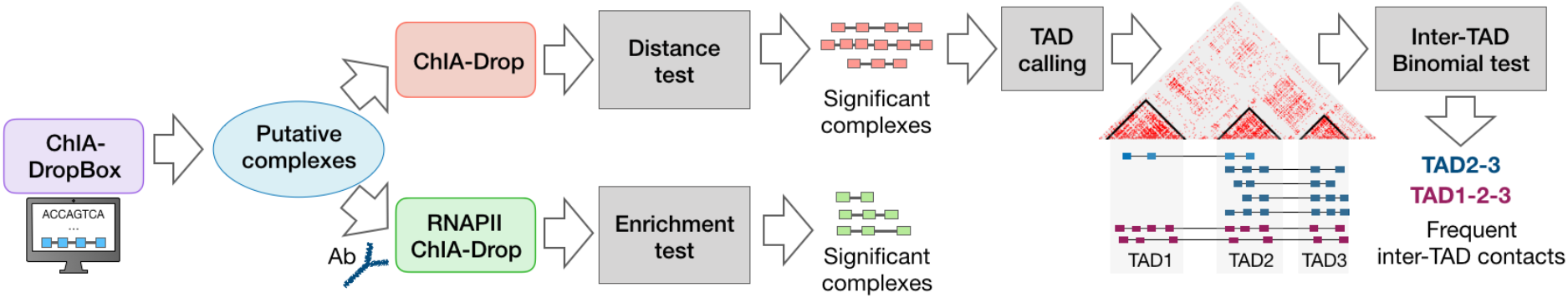
Summary of MIA-Sig algorithm. ChIA-Drop putative complexes from ChIA-DropBox pipeline are inputs to the distance test, which assigns p-values to each complex to quantify its significance. Refined complexes enable TAD calling directly on the multiplex data. A binomial test identifies frequent contacts among multiple TADs. RNAPII-enriched ChIA-Drop putative complexes are assigned significance according to the level of enrichment.

We envision that MIA-Sig will be broadly applicable to any type of multiplex chromatin interaction data ranging from ChIA-Drop, SPRITE, to GAM, under the aforementioned assumptions and with modifications (**Supplementary Note 1**). Here, we focused on the *Drosophila* ChIA-Drop and RNAPII ChIA-Drop data as a proof of concept and demonstrated that MIA-Sig filters and retains only the highly informative complexes. We anticipate that researchers will be able to utilize this refined data to dissect heterogeneous population within TADs and to identify interactions involving multiple regulatory elements.

## Data access

### Code availability

The MIA-Sig software is available at: https://github.com/TheJacksonLaboratory/mia-sig.git.

### Data availability

The ChIA-Drop (GSM6647523) and RNAPII ChIA-Drop (GSM3347525) data were downloaded from the Gene Expression Omnibus under SuperSeries accession number GSE109355. A link to the pure DNA ChIA-Drop data and processed files of relevant data are also available through the MIA-Sig github page.

## Acknowledgements

This study is supported by a Jackson Laboratory Director’s Innovation Fund (DIF19000-18-02). Y.R. is funded by 4DN (U54 DK107967) and ENCODE (UM1 HG009409) consortia. Y.R. is also funded by Human Frontier Science Program (RGP0039/2017), and supported by Florine Roux Endowment.

## Author contributions

M.K., M.Z., and Y.R. conceived the study. M.K. devised algorithms and wrote the MIA-Sig Python software with input from all authors. M.Z. developed and performed ChIA-Drop experiments. S.Z.T. developed and provided the ChIA-View software. S.Z.T., B.L., and D.C. contributed parts of the analyses. M.K., D.C., J.H.C., and Y.R. wrote the manuscript. All authors read and approved the final manuscript.

## Competing interests

The authors do not declare any competing interests.

## ONLINE METHODS

### Notation

An input dataset contains a set of chromatin complexes, each with 2 or more fragments. Let *OC_m_* be the set of fragments contained in the *m*-th ‘observed complex’ (OC), for *m* ∈ {1,2, …, *M*}, and *n* = |*OC_m_*| is the size of the set denoting the number of fragments in a complex. Each fragment *u* is subscripted by the complex index and superscripted by the fragment index, and encodes the genomic location of its origin expressed as a triplet of chromosome, start, and end position. The distance *d* between fragments 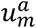 and 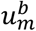 is 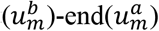, and neighboring (fragment-to-fragment; F2F) distances are encoded in a vector

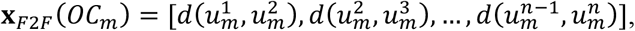

and the total distance is 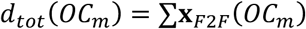; the probability vector 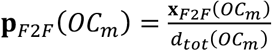. For example, if an 8-th complex 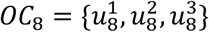 contains 3 fragments (chr2L, 100, 500), (chr2L, 1000, 1500), and (chr2L, 6000, 6500), then **x**_*F2F*_(*OC*_8_) = [500,4500], *d_tot_*(*OC*_8_) = 5000, and 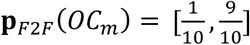. Finally, we can partition *M* complexes *OC*_0_, *OC*_2_, …, *OC_M_* into *F_j_*, where *j* is the number of fragments in a complex (*OC*_8_ belongs to *F*_3_ since it has 3 fragments).

### Distance test for non-enriched multiplex chromatin interactions data

#### Empirical null distribution and first distance test

Assuming that complexes are independent of chromosome, we perform the distance test separately for each chromosome. Motivated by the fact that each fragment class *F_j_* has distinct distributions in F2F distances, we construct the expected null background distribution by randomly rewiring fragments. Specifically, all neighboring distances **x**_*F2F*_(*OC_m_*) for *m* ∈ {1,2, …, *M*} are placed in a bucket *B*. For each observed *F_j_*, we randomly draw *j* − 1 elements (with replacement) from *B* to create 100,000 ‘expected complexes’ (EC) 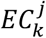 for *k* ∈ {1,2, …, 100000} and store them in 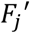. Note that since we only care about distance between fragments, we can assume that every fragment starts at (chr, 1, 500) and each fragment is of equal length. In practice, we store minimum information to save compute memory (implementation details below). For each *OC_m_* in *F_j_*, we compare its total F2F distance to total F2F distance in 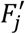 and record the proportion of expected complexes that have shorter distances than the observed complexes as the estimated ‘raw p-value’. Formally, for a *OC_m_* ∈ *F_j_*,

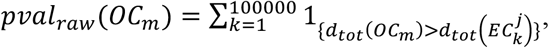

where 1_{*}_ is an indicator function. Assuming that complexes in each fragment class are independent, we subsequently separate the raw p-values by *F_j_* and adjust them for multiple hypothesis testing using Benjamini-Hochberg method (Benjamini, Hochberg, 1995) with false discovery rate (FDR) of 0.1. The complexes with adjusted p-value ≤ 0.1 are considered to be statistically significant and are classified as ‘pass1’ (*F*_*j,pass*1_). Of those insignificant complexes with adjusted p-value > 0.1, we ‘fail 1’ (*F*_*j,fail*1_) those with 2 fragments (*OC_m_* ∈ *F*_2_ with adjusted p-value > 0.1) and treat others in a separate category called ‘defer’ (*F_j,def_*). These ‘deferred’ complexes are passed onto the entropy filter to correct for droplet contamination.

#### Entropy filter

Some complexes in the ‘deferred’ category may be due to the experimental noise that can be computationally detected. Specifically, this step aims to computationally correct for the undesired phenomenon of a droplet containing more than one chromatin complex (referred to as ‘doublet’ for two, ‘triplet’ for three, and ‘multiplet’ for 2 or more). In single-cell RNA-seq (scRNA-seq; single-cell transcriptome) experiments, the outcome of a doublet would be a vector of real numbers indicating average expression of the two cells. By contrast, ChIA-Drop data only provide binary values indicating if a fragment was captured or not, with a variable number of fragments. Therefore, the effect of two complexes accidentally being encapsulated in a single droplet would be a large distance in the data. This assumption is based on the observation from Hi-C and ChIA-PET data analysis that true interactions occur within certain range of genomic span. Our goal is to identify complexes with one dominating distance between fragments. Using the probability vector of the neighboring distance, we quantify the likelihood of a dominating event. Formally, for an observed complex *OC_m_* with *n* fragments and **p**_*F2F*_(*OC_m_*) = [*p*_1_,*p*_2_, …, *p*_*n*−1_], we compute the normalized Shannon entropy (Shannon, 1948)

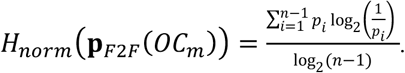

The normalization factor log_2_(*n* − 1) ensures that *H_norm_*(***x***) ∈ [0,1] for any probability vector ***x***. Generally, *H_norm_* is small when only one or two of *p_i_*of is large, in which case we presume that a complex is a multiplet and need to separate into singlets. For each observed complexes in the ‘deferred’ category, we compare its normalized Shannon entropy to the average normalized Shannon entropy of the expected complexes in the corresponding class; if the former is smaller, then we separate the observed complex at the longest distance interaction. In other words, for *OC_m_* ∈ *F_j,def_*, if

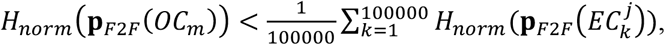

then *OC_m_* is separated into

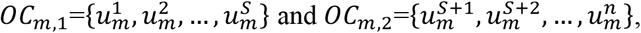

where 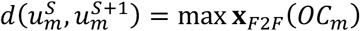. Furthermore, if the second largest distance is at least 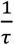 of the largest distance, we also separate at the second longest distance. *τ* is a variable parameter and we set it to 2 in our analyses; the larger the *τ*, the more likelihood of a ‘second cut’ (implying a triplet). The resulting sub-complexes are placed in *F_J,def,filt_*, and are now subject to the second distance test. Note that we did not perform any statistical test in this step; only performed filtering. Also, the Shannon entropy merely serves as a quantification measure for a single complex and should not be confused with the heterogeneity of all complexes in the ChIA-Drop data.

#### Second distance test

We repeat the distance test after correcting for possible doublets and triplets. For a *OC_m,∗_* ∈ *F_j,def,filt_*,

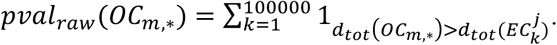

We adjust raw p-values using Benjamini-Hochberg method with false discovery rate (FDR) of 0.1. The complexes with adjusted p-value ≤ 0.1 are classified as ‘pass2’ (*F*_*j,pass*2_) and others are ‘fail2’ (*F*_*j,fail*2_). A diagram of the distance test is illustrated in **Supplementary Figure 1a**.

#### Implementation, results, and analysis

MIA-Sig takes putative chromatin complexes as the input, which are results of the ChIA-DropBox (Tian et al., 2019) data processing and visualization pipeline. The ‘distance test’ python (v3.6) script encompasses all parts using the following packages: numpy, random, statsmodels, itertools, os, sys. We used the parameters --gen dm3 --fdr 0.1 --cef 2 --sz 100000 to run the script on GSM3347523 dataset, which used 1.8 Gigabyte of memory and 13 minutes of cpu time. To save memory, we store minimal information for the null: total distance for expected complexes, and their mean entropy for each fragment class. Two runs with the same parameters should yield identical results because we set seeds in the construction step for the expected complexes. By saving the first 1000 expected complexes for each class in a chromosome, we can compare our expected null model to the biological null model, which is the ‘pure DNA’ described in (Zheng et al., 2019). Plotting the neighboring distances, we observed that both the computational null and pure DNA are unimodal with peaks between 1 Mbps and 10 Mbps for all classes (**Supplementary Figure 1b**). After confirming that our expected complexes do emulate long-range noise, we obtained detailed statistics of each step resulting in 55,995 significant complexes (**Supplementary Figure 1c**). Complexes in each of the ‘original’, ‘significant’ (‘pass1’+’pass2’), and ‘insignificant’ (‘fail 1’+ ‘fail2’) categories are converted into a .short format by enumerating over all pairs of fragments in a complex. Three .short files are then converted into .hic files via juicer (v1.7.5) to be visualized in juicebox. A 5 Mbps window on chr3L shows that the original data exhibit both the signal and noise, which are separated by MIA-Sig into significant and insignificant, respectively (**Supplementary Figure 2a**). The original observed complexes have a bimodal distribution for high fragment classes, which is a distinct behavior from the null distribution (**Supplementary Figures 1b, 2b**). The density plot further support that significant complexes retained short distances or a mix of short and long distances. By contrast, insignificant complexes are only comprised of unimodal long distances (**Supplementary Figure 2b**). Consistent with an observation that high-fragment complexes contribute to the structure more than the low-fragment complexes (Zheng et al., 2019), MIA-Sig assessed the majority of high-fragment complexes as significant (**Supplementary Figure 2c**). We next investigated the effects of the entropy filter, which was designed to remove doublets and triplets. Of the 1,452,878 complexes in the deferred category ranging from *n* = 3 to *n* = 8, MIA-Sig identified 60% (869,065) to be singlets, 34% (498,291) to be doublets, and 6% (85,522) to be triplets, yielding 548,342 singletons (*F*_1_) and 1,573,871 complexes (*F*_≥2_) (**Supplementary Figure 3**). For each class, singletons had the highest normalized Shannon entropy, followed by doublets and triplets. The entropy filter step allowed MIA-Sig to identify additional 15,055 complexes as significant, which amounts to 27% of the total significant complexes.

### TAD calling for non-enriched multiplex chromatin interactions data

#### Generating 1D signal track

Existing TAD calling algorithms for pairwise Hi-C data generally fall into two categories: 1) signal segmentation after conversion from 2D contact maps into 1D tracks measuring interaction intensities along the genome, 2) community detection directly on the 2D heatmap by treating each bin as a node on an undirected graph. We take the first approach and convert our complexes into 1D signal track. A conventional pairwise approach would enumerate over all pairs of fragments in a complex and record their spans. However, multi-fragment complexes may over-contribute since the number of pairs grows quadratically: 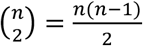, where *n* is the number of fragments in a complex. Instead, we allow each complex to only contribute linearly in *n* by recording its span weighted by *n*. More precisely, coordinates are (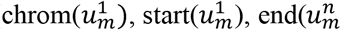, *n*) for an *OC_m_* with *n* fragments. We finally obtain a ‘weighted complex span coverage’ by accumulating the coordinates over all given complexes.

#### Smoothing and segmentation

Our next task is to segment the 1D track into regions with high signal and annotate them as TADs. In an ideal case, we can achieve this goal by computing the slope of the signal **s** and by recording critical points where the slope is 0. However, our signal has a basepair resolution and thus is not smooth, resulting in too many false critical points. A common way to smooth the signal is by a moving average window, but using a large window size would lose the resolution and yield TADs with fuzzy boundaries. Moreover, due to the inherent nature of TAD sizes, a window size parameter optimal in one region may not be optimal in another. We avoid this parameter tuning step by instead applying a discrete wavelet transformation, which decomposes signal into high-frequency component and low-frequency component (Mallat, 1989). Of note, the low-frequency component generally retains the smoothed version of the signal without affecting the shape, which is helpful for us to find accurate TAD boundaries (**Supplementary Figure 4**). Using this ‘smoothed’ signal, we compute the slope and fine-tune TAD coordinates.

#### Implementation, results, and analysis

The ‘tad calling’ python (v3.6) script encompasses all parts using the following packages: numpy, os, scipy, pywt, itertools, sys. We used the parameters --cat PASS --bs 1000 –sp drosophila --r dm3 to run the script on significant complexes from the distance test of GSM3347523 dataset, which used 84 Megabyte of memory and 1 minute of cpu time. Before generating the 1D signal track, we separate two fragments if they are more than 100 Kb apart, based on the upper range of general TAD sizes by organisms (Dekker and Heard, 2015). Coverage was generated by BEDtools (Quinlan et al., 2010) using ‘genomecov’ function and the coverage is binned into 1 kb windows via ‘makewindows’ and ‘map’ commands. Signal smoothing was done by pywt package using the parameters ‘bior1.1’ for wavelet function and ‘3’ for the level. MIA-Sig called 335 TADs over the 6 chromosomes, with a median size of 200 Kb (**Supplementary Figure 5a**). For a comparison, we also tested insulation score as follows: .hic file (of all pairs of fragments) are converted into contact matrices via Juicer’s ‘dump’ function with a dense matrix option (-d) in the Juicer tool (v1.7.5); insulation score script (https://github.com/dekkerlab/cworld-dekker/tree/master/scripts/perl) is executed with 100Kb insulation square size, 100Kb delta window size for 10Kb resolution contact maps with balanced normalization. Insulation score (InS) called 513 TADs with a median of 150Kb, and did not call any TADs larger than 500 Kb (**Supplementary Figure 5b,c**). When we examined the gaps (defined as regions between two TADs, if any), MIA-Sig also had wider size range than InS (**Supplementary Figure 5d,e**). For each TAD called by MIA-Sig and InS, we compute the total H3K27me3 signal and plot the genome-wide behavior (**Supplementary Figure 6a**). Overall, MIA-Sig has higher inactive signal in TADs than InS. The gap regions in *Drosophila* are known to be transcriptionally active and should positively correlate with H3K27ac signal. We confirm that MIA-Sig has slightly higher median active signal than InS (**Supplementary Figure 6b**). Note that we did not perform any normalization by region size because both algorithms segment the genome into either a TAD or a gap, so the region size should also be a feature. Histone marks provide biological evidence that MIA-Sig TADs are inactive and gaps are active, but ChIA-Drop fragment counts provide a direct measure of TAD and gap intensities. Using the BEDtools command ‘intersect -c’, we count the number of fragments in each region. MIA-Sig generally captured more fragments in TADs than InS did (**Supplementary Figure 6c**), and less fragments in gaps than InS (**Supplementary Figure 6d**). Finally, we annotate each fragment in significant and insignificant complexes as ‘TAD’ or ‘gap’ as called by MIA-Sig. For each complex, we count the number of TADs with at least 2 fragments within each TAD. Only 5% of the insignificant complexes had fragments in 1 or 2 TADs, and the rest were not contributing to the TAD structure (**Supplementary Figure 7a**), validating the observation from 2D heatmaps. By contrast, only 26% of the significant complexes were not in TADs, a majority (51.3%) in intra-TAD interactions, and many (23%) connected two or more TADs. By observing that 12,884 complexes involve 2 to 21 TADs, we next sought to characterize if multiple complexes connect the same set of TADs

### Inter-TAD binomial test for non-enriched multiplex chromatin interaction data

#### Motivation and intuition

Our goal is to evaluate the statistical significance of these TAD combinations based on the frequency of occurrence measured by the number of complexes therein. The problem is simple for a pair of TADs: we may treat a TAD as a ChIA-PET loop anchor and apply tools based on hypergeometric test. However, our data are now multi-dimensional. For instance, suppose that there are 5 TADs, and 5 combinations ‘A-C’, ‘B-C’, ‘B-C-D’, ‘A-B-E’, ‘A-D-E’ (**Supplementary Figure 7b**). The pair ‘B-C’ appears 4 times on its own, but also appears 3 times as a part of the triple ‘B-C-D’. Moreover, some parts of a combination may appear elsewhere with the same number of TADs: given ‘B-C-D’ and ‘A-C-D’, ‘C-D’ appears twice. Therefore, we propose a counting scheme based on occurrence of ‘expanded pairs’.

#### Methods

The notations used defined in this section are independent from those in other sections. We let the ¿th combination be 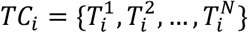, where each 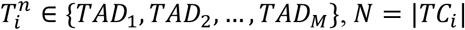 is the number of TADs involved, and we partition each *TC_i_* into the same class *G_j_* if |*TC_i_*| = *j*. All pairs of TADs in *TC_i_* are in *Pa*(*TC_i_*) = {{*r, s*}: *r* ≠ *s*, for *r*, *s* ∈ *TC_i_*} and 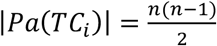. For each *TC_i_*, we record the number of pairs in the same class as

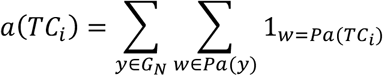

and the number of exact appearance in higher class as

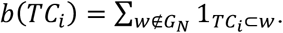

Using these two numbers, we compute the appearance of ‘pairs’ in same class and higher class:

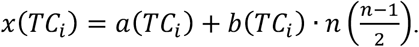

Finally, we perform the binomial test with *x*(*TC_i_*) as the number of success, 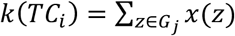 as the number of trials, the probability of success hypothesized as 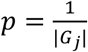; the alternative hypothesis is that the observed probability is greater than the expected probability *p*. A detailed example is provided using same notations (**Supplementary Figure 7b**).

#### Implementation, results, and analysis

A python script ‘inter-TAD binomial test’ implements the method using packages numpy, itertools, scipy, statsmodels, os, and sys. Of 6,861 unique combinations involving 2 to 21 TADs, 915 (13%) were identified as statistically significant. An example illustrates that a pair of TADs with strong signal in the heatmap and many complexes in the linear view has lower p-value than that with weak signal (**Supplementary Figure 7c**). Here, we assumed that frequency of interactions between TADs is independent from their distance and sizes, and we also did not distinguish contacts with 2 fragments from those with 10 fragments. These parameters may be incorporated in the future version.

### Enrichment test for RNAPII-enriched multiplex chromatin interaction data

#### Motivation

The above sections are designed to analyze non-specific multiplex interaction data analogous to the Hi-C data. With an additional step of chromatin immunoprecipitation, protein-enriched multiplex data reveal protein-specific interactions similar to the population average ChIA-PET loops. In a typical ChIA-PET analysis, loops anchored in strong binding peaks are considered to be more reliable than those with weak or no peaks. Extending this notion to the multiplex data, we developed an enrichment test for RNAPII ChIA-Drop data. Our end goal is to retain complexes with fragments in strong binding peaks. One approach is to call peaks and only keep complexes that overlap the peak regions. However, peak calling algorithms have their own model assumptions that may not hold for ChIA-Drop data. Even with accurate peak regions, assigning statistical significance to each complex is not a trivial problem since the null distribution is unclear. Thus, we take an alternative—inevitably the computationally expensive— approach by sampling the background null distribution for each complex.

#### Statistical test

The idea is to take the observed complex and place it on a random location of the same chromosome and compare the mean coverage between the observed and the expected. Through many rounds of re-sampling, we obtain the p-value by counting the number of occurrences in which the expected coverage exceeds the observed coverage (**Supplementary Figure 8a**). More precisely, for an observed complex 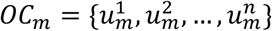, we randomly draw an integer 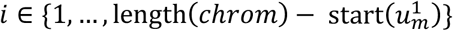 and the 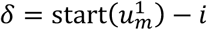. The first expected complex is then 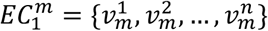, where 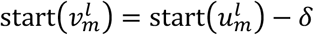, and 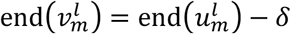 for all *l* ∈ {1, …, *n*}. Repeating this process 10,000 times, we obtain 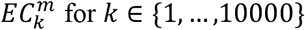. We can then compute the raw p-value of the mth observed complex as:

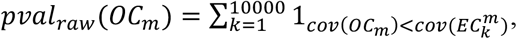

where 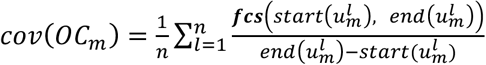 and ***fcs***(*x,y*) is the mean ‘fragment coverage signal’ between coordinates *x* and *y*. Raw p-values are separated by chromosomes and are adjusted via the Benjamini-Hochberg method with false discovery rate (FDR) of 0.1. The complexes with adjusted p-value ≤0.1 are considered to be statistically significant and are classified as ‘pass’; others are considered insignificant or ‘fail’.

#### Implementation, results, and analysis

A python script ‘enrichment test’ utilizes packages numpy, random, statsmodels, os, sys. GSM3347525 RNAPII ChIA-Drop data are pre-processed to exclude fragments mapped to the repetitive regions in the genome (dm3.rmsk.bed), and 769,803 complexes remain as ‘GSM3347525NR’. The most time-consuming part of the algorithm is to obtain the fragment coverage at a given location, since we need to search for a start and end index in a bedgraph or a bigwig file. With at least 769803 * 2 * 10000 = 1.54 × 10^10^ operations, we realized that python implementations of exact search would be intractable. As means to reduce the runtime, we store the bedgraph file into bins of size 10 bps and store only the 4^th^ column ‘value’. The solution then turns into a simple lookup operation, yielding an approximation that is close to the exact solution. Our code is ‘parallelized’ by chromosome, each using around 5 hours cpu time and 230 Megabytes of memory (**Supplementary Figure 8b**). MIA-Sig identified 190,226 complexes (24.7%) as statistically significant. We ensure that our empirical null distribution does behave randomly by comparing the enrichment scores of the observed complexes in chr2L with those of 1,000 expected complexes generated for each observed complex (**Supplementary Figure 8c**). Zooming in further, we note that the histogram of the observed is shifted to the right of the histogram of the expected null (**Supplementary Figure 8d**). Using the active and inactive regions defined in (Zheng et al., 2019), we count the number of fragments therein for significant and insignificant complexes (**Supplementary Figure 9a**). For each active and inactive region, we compute the number of significant complexes fragments and their log10 values are plotted (**Supplementary Figure 9b**); K-S test supports that significant complexes are indeed more likely to be in active regions than inactive regions. By contrast, insignificant complexes have no bias towards or against active regions (**Supplementary Figure 9c**). We define a gene promoter as +/− 1kb from the transcriptional start site (TSS) annotated by UCSC genome browser. Note that typically +/− 250 bp is used for drosophila, but we extend it to accommodate ChIA-Drop protocol-specific features. A gene is active (6466 genes) if total RNA-seq level is greater than 5, and inactive (8874 genes) otherwise. A fragment is ‘active promoter’ if it overlaps the promoter of an active gene. In general, significant complexes have higher proportion of promoter fragments than insignificant complexes (**Supplementary Figure 9d**), and the skew is more pronounced for active promoters (**Supplementary Figure 9e**). Inactive promoters serve as a control, in which both significant and insignificant complexes display similar patterns in the number of inactive promoter fragments (**Supplementary Figure 9f,g**).

## Supplementary Figures and legends

**Supplementary Figure 1:**
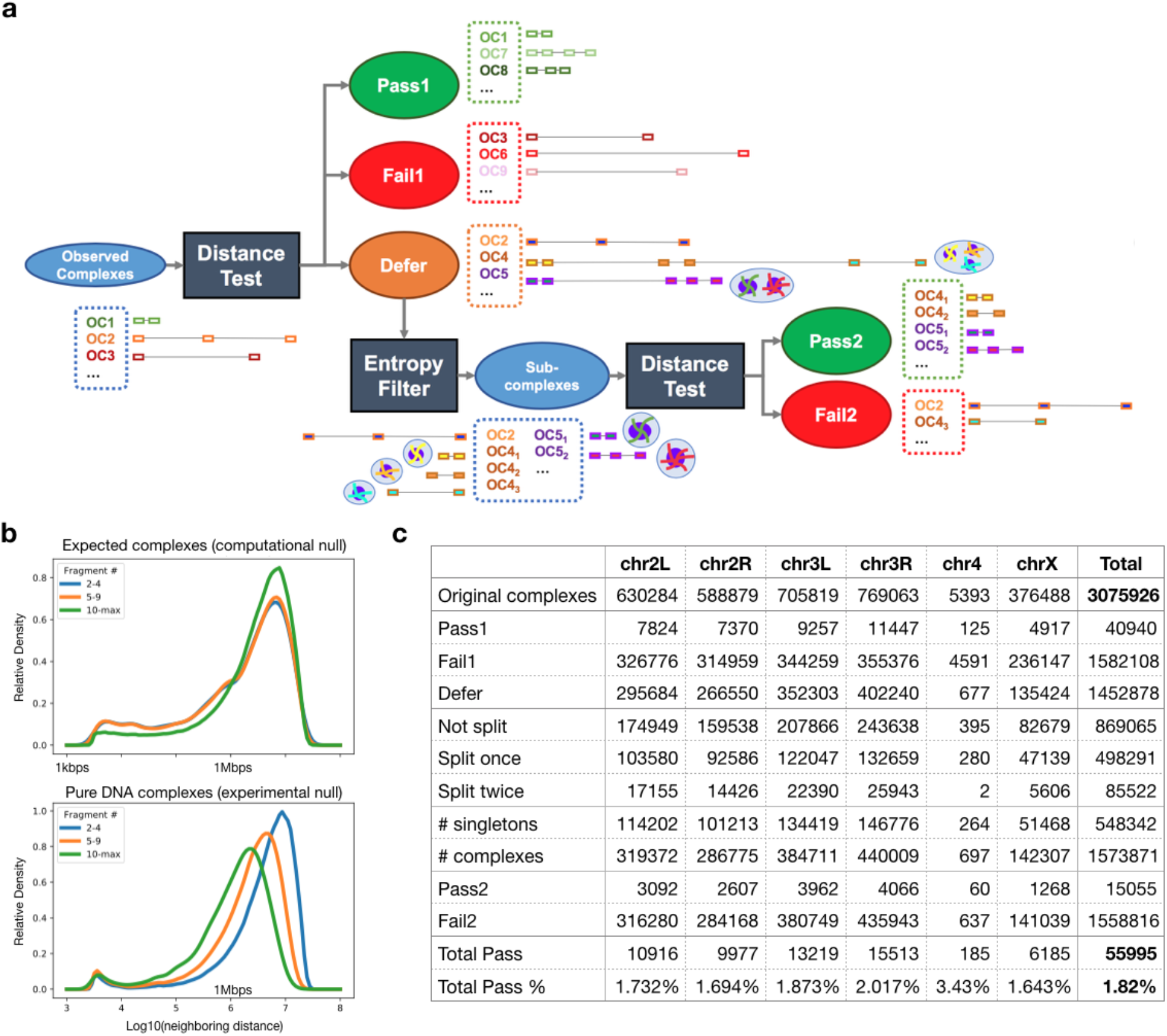
Overview of distance test, comparison of computational and experimental null distribution, and summary statistics. (a) A diagram of distance test encompasses both the statistical test and the filter for correcting multiplets, which is defined as droplets with more than one chromatin complex. All observed complexes get tested for distance, and those with significant p-values ‘pass’; of those with insignificant p-values, 2-fragment complexes ‘fail’, and others are ‘deferred’. Entropy filter separates multiplets into singlets and resulting complexes are subject to the second distance test. (b) Log10 of neighboring fragment-to-fragment distances of expected complexes and pure DNA complexes are computed by fragment number class. Density plot illustrate that both datasets comprise mostly large distances. (c) A comprehensive statistics of complexes at each step of the distance test. A singleton is a complex with only one fragment, and complex has more than one fragment.

**Supplementary Figure 2:**
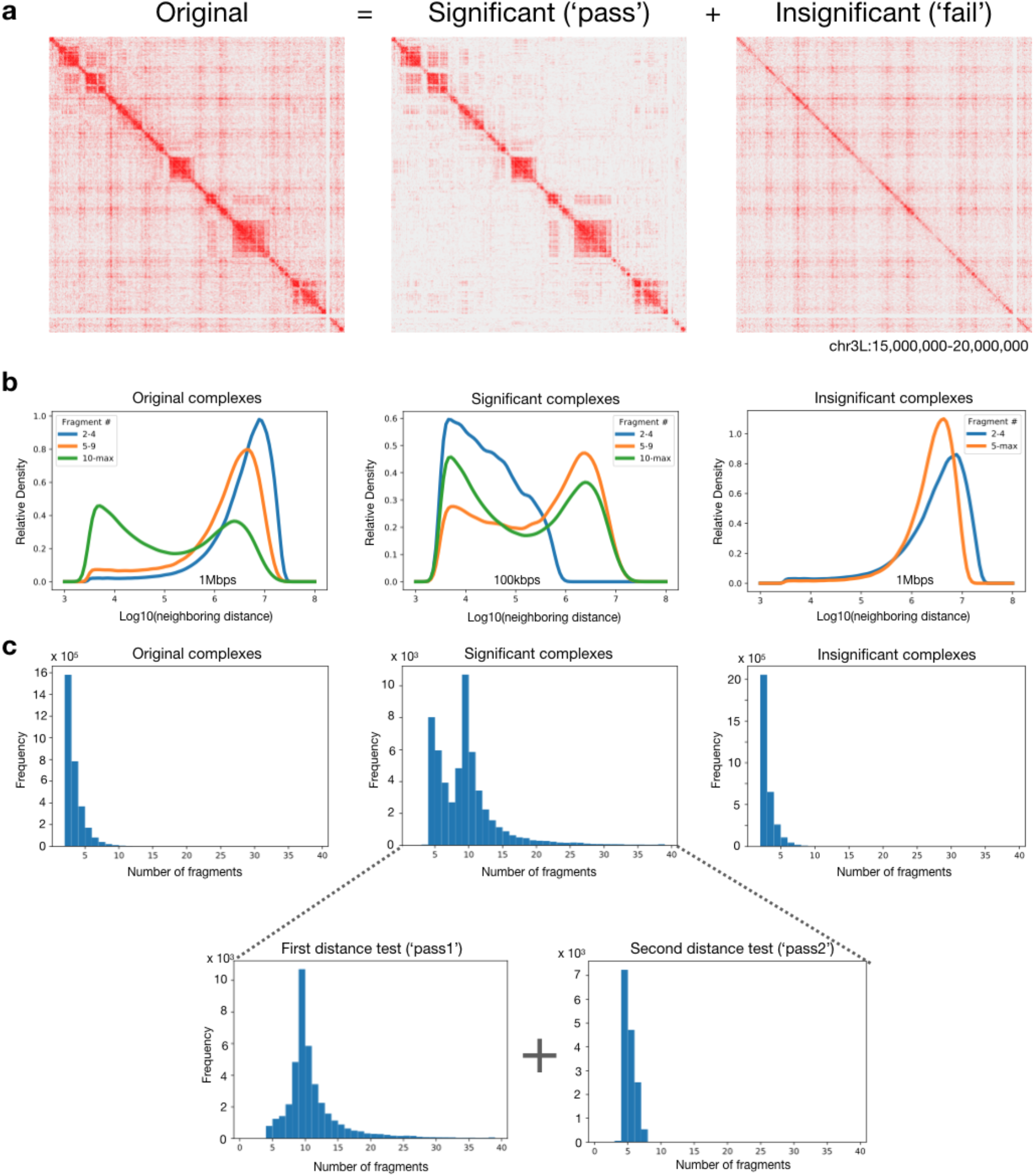
Characteristics of original, significant, and insignificant complexes. (a) A 2D heatmap is shown for each of original, significant, and insignificant complexes after the distance test. Significant complexes contribute to the TAD structures, and insignificant complexes have seemingly random behavior throughout the heatmap. (b) Neighboring distances are plotted as in Supplementary Figure 1b. Only high-fragment original complexes are bimodal. Significant complexes are either bimodal or unimodal with distances less than 1 Mbps; majority of insignificant complexes contain large distances. (c) A histogram of the number of fragments in each complex indicates that complexes with many fragments pass the first distance test, and additional complexes with 4 to 8 fragments pass the second test after entropy filter. Insignificant complexes are mainly low-fragment complexes.

**Supplementary Figure 3:**
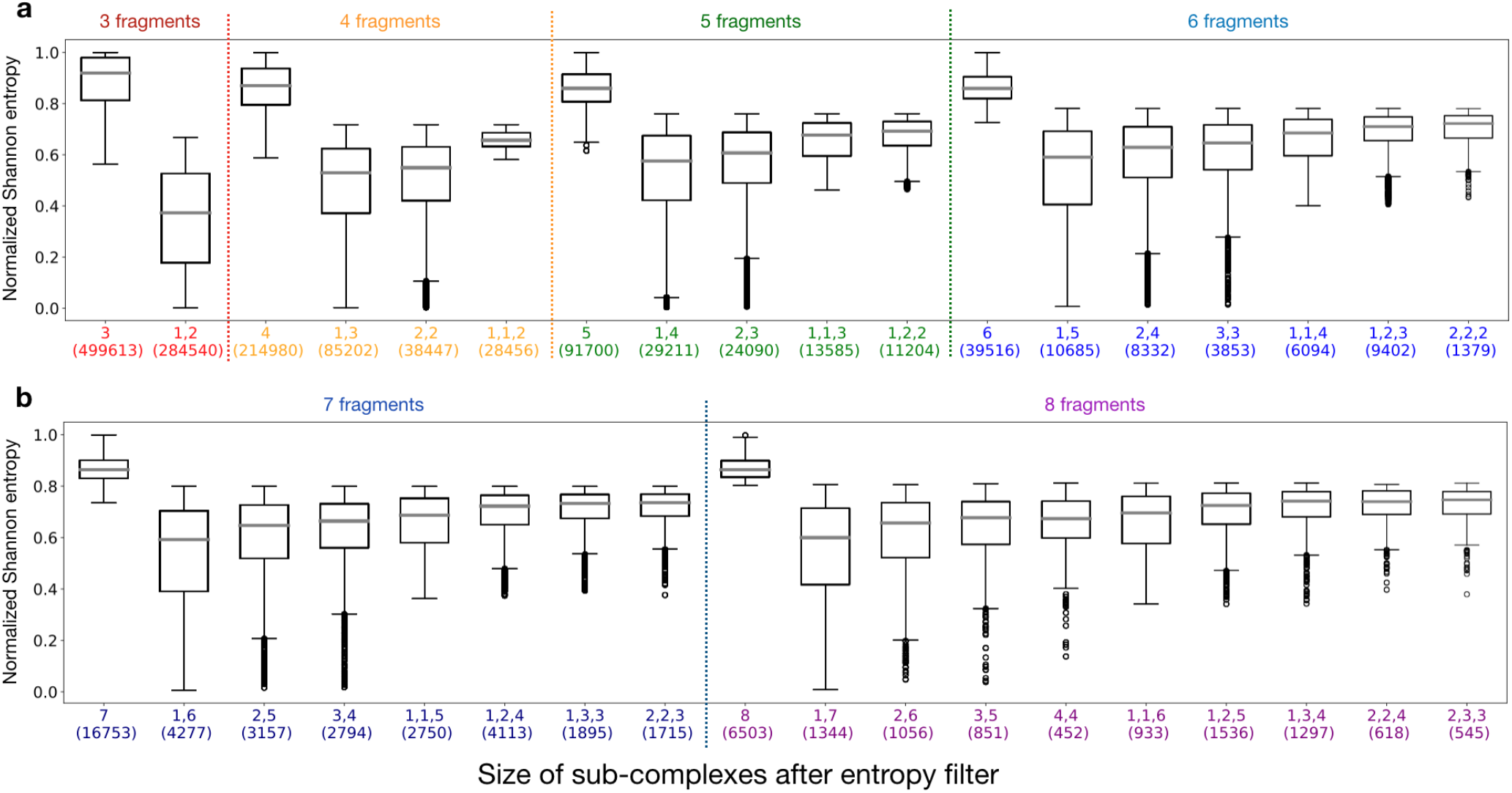
Effects of the entropy filter. (a) A box plot of normalized Shannon entropy is provided for sub-complexes after filter. Of complexes with three fragments (in *F*_3_), 499,613 are identified as ‘singlets’ due to high entropy, and 284,540 are considered to be ‘doublets’ due to low entropy. x-axis is labled with the size of sub-complexes, with counts in a parenthesis. For instance, a 5-fragment complex can be split into a singleton (1) and a 4-fragment complex (4), or into a complex (2) and another complex (3); alternatively, a triplet has three sub-complexes (1,1,3 and 1,2,2). A general trend is that entropy is highest for those without any splits, lowest for a doublet with a singleton, and increases as the size of sub-complexes balance to be roughly equal. (b) Complexes in *F*_7_ and *F*_8_ are plotted in an identical procedure as part (a).

**Supplementary Figure 4:**
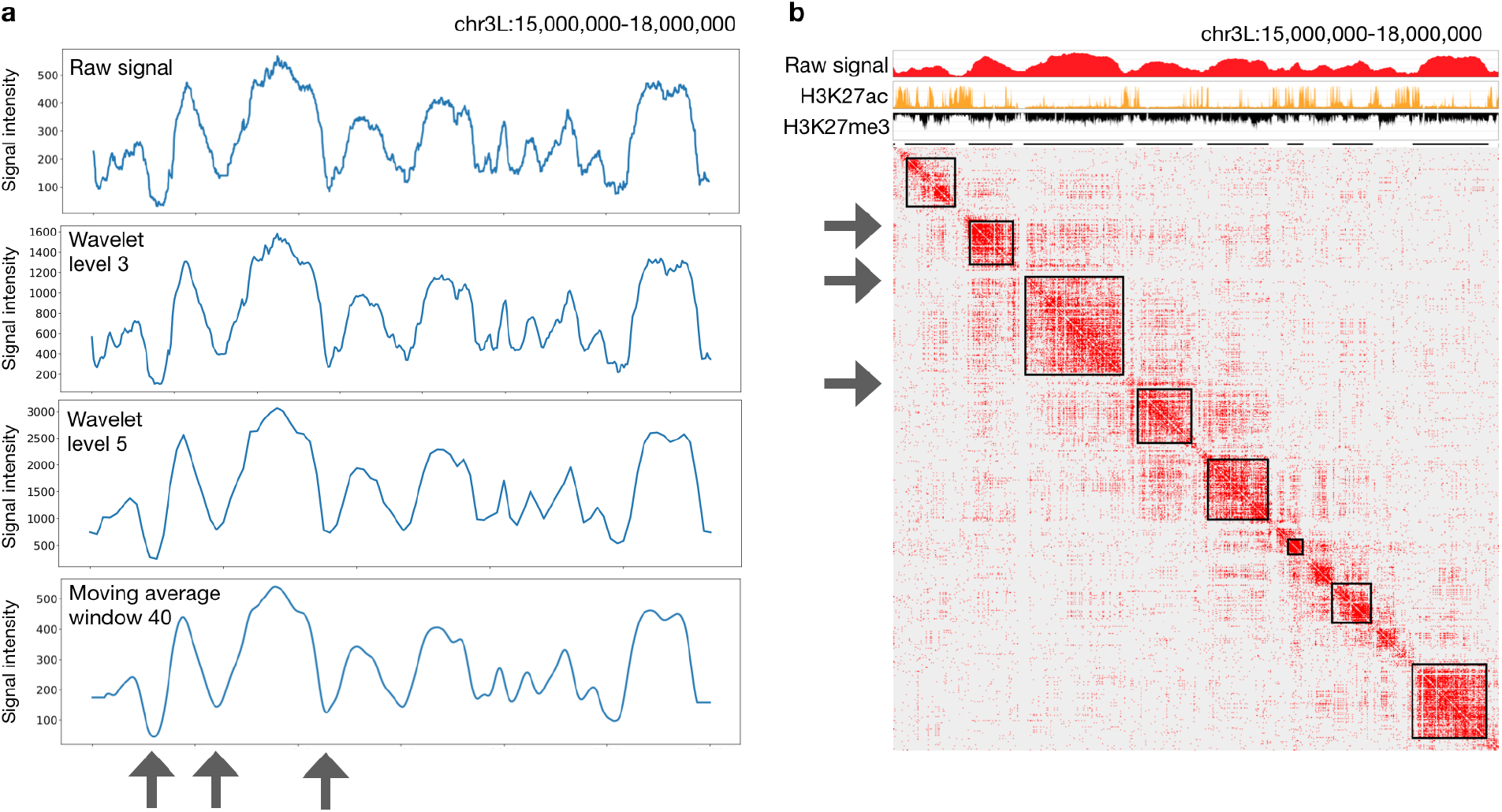
Ideas behind the MIA-Sig TAD calling algorithm. (a) A raw signal track is generated from significant complex span with preprocessing. Wavelet transformation of the raw signal with level 3 yields a moderately smoothed version, and that with level 5 removes more noise (both using biorthogonal wavelet). Similar results are obtained with a triangle moving average window of size 40 bps, but ‘valleys’ are narrower than those by wavelet method. (b) MIA-Sig distinguishes clear gaps between TADs (indicated by three arrows) possibly due to wavelet method.

**Supplementary Figure 5:**
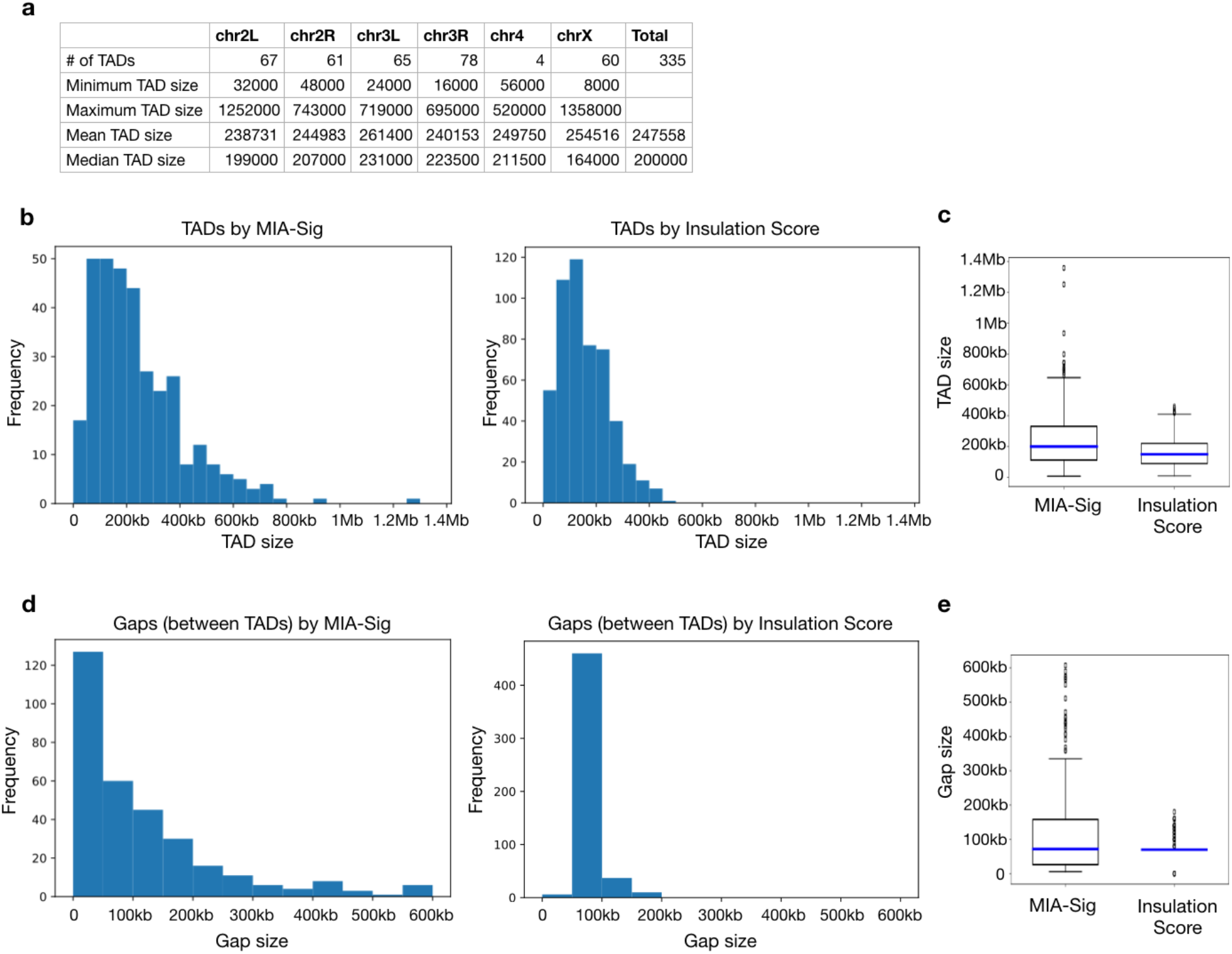
Statistics of TAD and gap sizes called by MIA-Sig and Insulation Score. (a) Statistics of 335 TADs called by MIA-Sig for each chromosome. (b) Histogram of sizes of TADs called by MIA-Sig (left; n=335) and Insulation Score (right; n=513). (c) Boxplot of TADs called by MIA-Sig (median=200kbps) and Insulation Score (median=150 kbps) (Mann-Whitney U test p-value= 8.16 × 10^-13^). (d) Histogram of sizes of gaps between TADs by MIA-Sig (left; n=318) and Insulation Score (right; n=513). (e) Boxplots of gaps by MIA-Sig (median=72 kbps) and Insulation Score (median=70kbps) (Mann-Whitney U test p-value= 0.29).

**Supplementary Figure 6:**
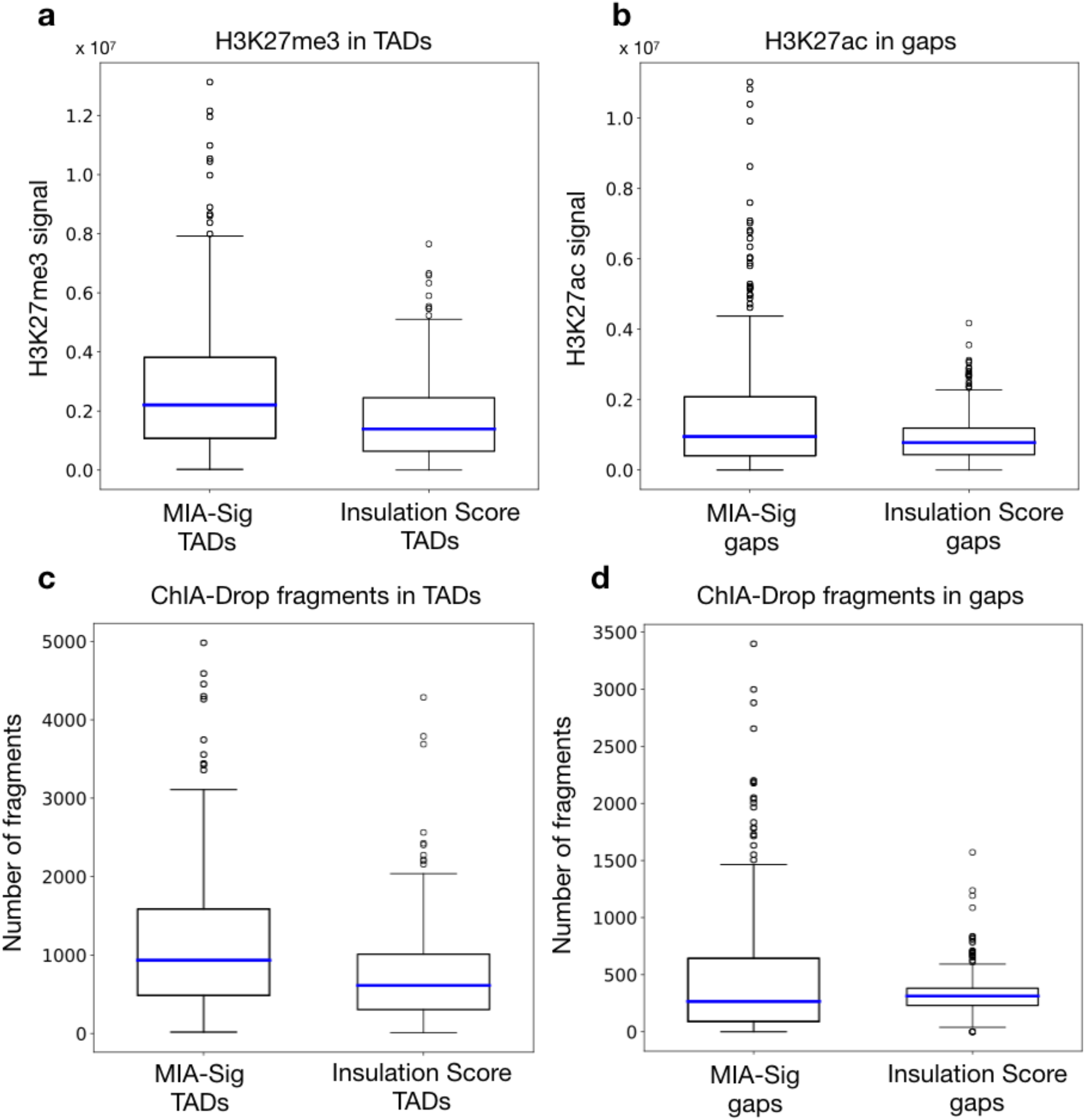
Comparison of TADs and gaps by MIA-Sig and Insulation Score. (a) Total H3K27me3 signal is plotted for each TAD called by MIA-Sig (median= 2.2 × 10^6^) and Insulation Score (median= 1.4 × 10^6^) (one-sided Mann-Whitney U test p-value= 7.33 × 10^-12^). (b) Total H3K27ac signal is plotted for each gap defined by MIA-Sig (median= 9.5 × 10^5^) and Insulation Score (median= 7.8 × 10^5^) (one-sided Mann-Whitney U test p-value= 0.0003). (c) Number of fragments from significant ChIA-Drop complexes are recorded for each TAD. MIA-Sig TADs have more fragments (median=933) than Insulation Score TADs (median=613) (onesided Mann-Whitney U test p-value= 1.5 × 10^-13^). (d) In gap regions between TADs, MIA-Sig has less fragments (median=266.5) than Insulation Score (median=313), though not statistically significant (one-sided Mann-Whitney U test p-value= 0.1).

**Supplementary Figure 7:**
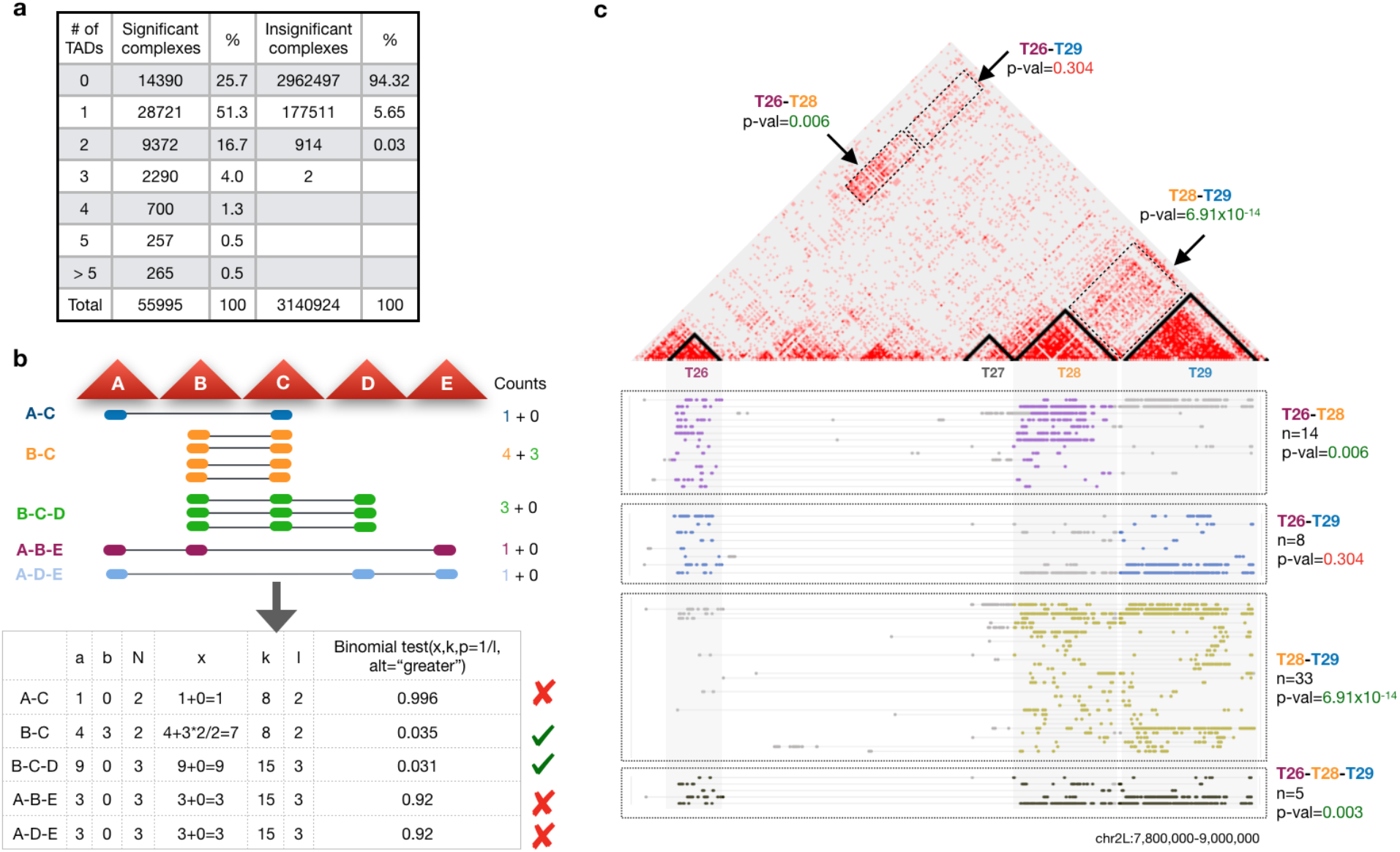
Inter-TAD binomial test. (a) For each complex, a TAD interaction is valid if two or more fragments are enclosed in TADs called by MIA-Sig. Most (51.3%) of significant complexes are intra-TAD interactions, yet 23% involve 2 or more TADs. By contrast, the majority (94.32%) of insignificant complexes are not in any TAD. (b) Inter-TAD binomial test is designed to assess significance of the 23% identified in part (a). Given a set of complexes spanning over 5 TADs ‘A’, ‘B’, ‘C’, ‘D’, and ‘E’, we count the number of complexes in a specific TAD combination and use the counts to test for significance via one-sided binomial test (details in the Method). Only ‘B-C’ and ‘B-C-D’ are identified as significant contacts. (c) An example of complexes involving 3 TADs with identifications ‘T26’, ‘T28’, and ‘T29’. ‘T26-T28’, ‘T28-T29’ and T26-T28-T29’ are classified as significant contacts, whereas ‘T26-T29’ is covered by only 8 complexes and thus insignificant.

**Supplementary Figure 8:**
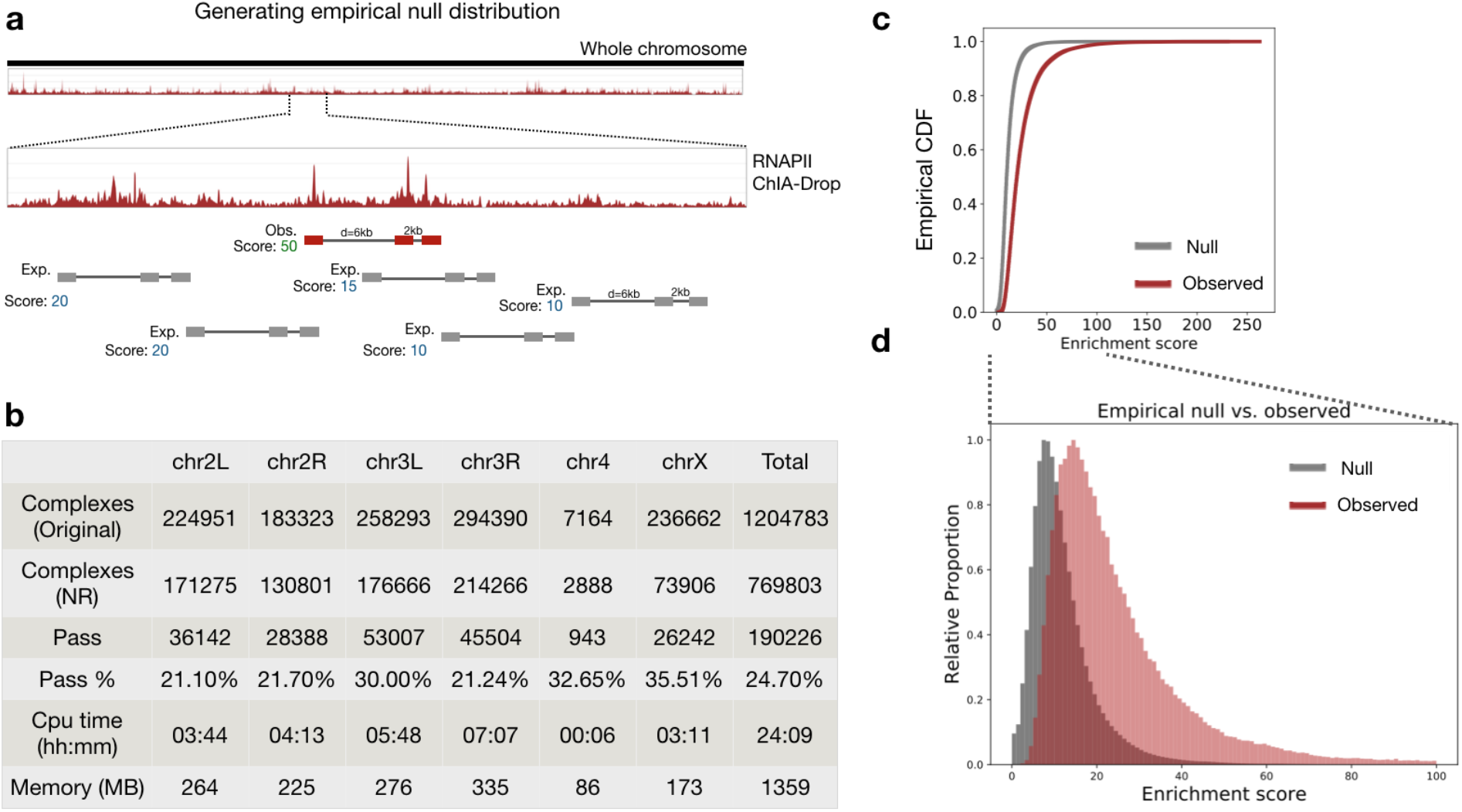
Overview of enrichment test for RNAPII ChIA-Drop data. (a) Empirical null distributions are generated by taking the observed complex and placing it in a random location of the same chromosome; ‘enrichment score’ is computed as the mean fragment coverage. (b) Summary statistics of complexes after enrichment test. (c) Empirical cumulative distribution function of the ‘enrichment score’ is plotted for expected (computational null) and observed complexes (one-sided Kolmogorov-Smirnov test statistic=0.47, p-value< 2.2 × 10^-16^). (d) A closer view of the distribution reveals that enrichment scores of observed complexes are shifted to the right of expected (null) complexes.

**Supplementary Figure 9:**
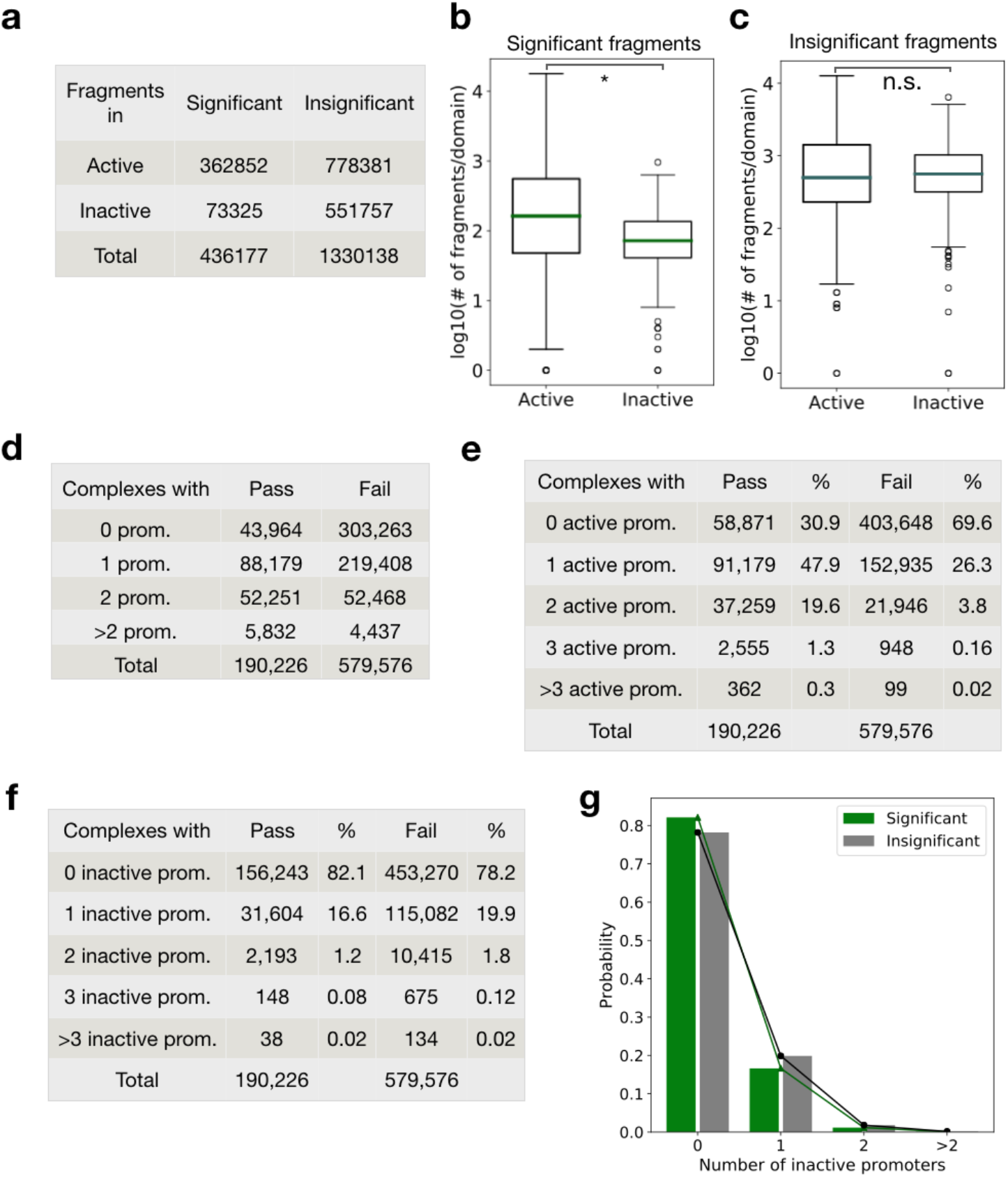
Comparison of significant and insignificant RNAPII complexes. (a) A majority (83%) of the fragments in significant complexes is in active region (cf. 58% of insignificant complexes). (b) For each active and inactive domain, number of fragments of significant complexes are plotted. Genome-wide, significant fragments are more likely to be in active regions than inactive regions (‘*’ denotes significant; Mann-Whitney U test p-value= 3.7 × 10^-22^). (c) Insignificant fragments have no preference over active and inactive regions (‘n.s.’ denotes not significant; Mann-Whitney U test p-value= 0.23). (d) Number of complexes with 0, 1, 2, more than 2 promoter fragments are recorded for significant (‘Pass’) and insignificant (‘Fail’) complexes. (e) Most (69%) of the significant complexes have at least one active promoter fragment (cf. 30.4% of insignificant complexes). (f) In both significant and insignificant complexes, around 80% had no fragment annotated as an inactive promoter. (g) Distribution of inactive promoters have similar pattern for significant and insignificant complexes.

**Supplementary Figure 10:**
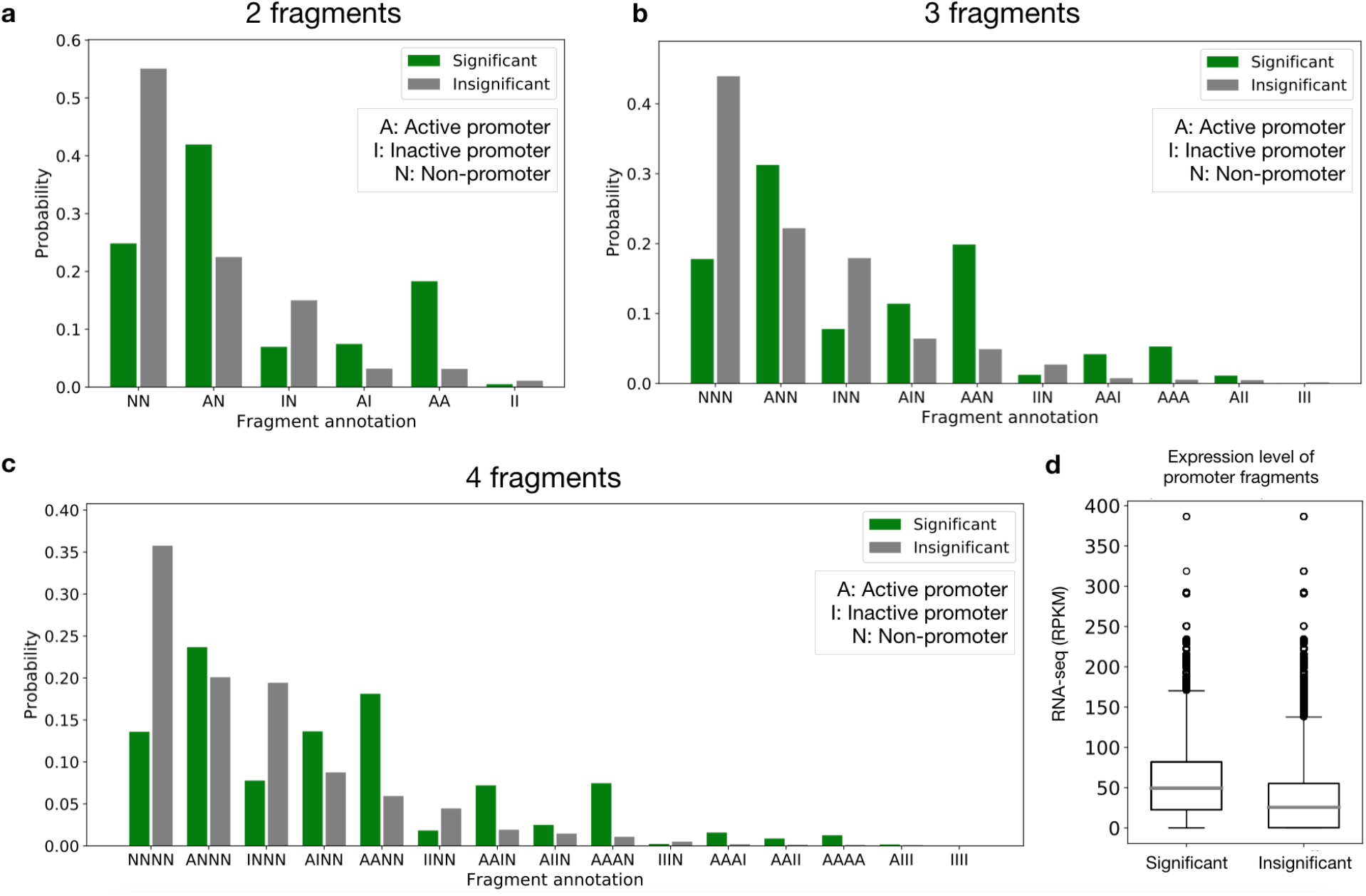
Annotation of fragment as an active promoter, inactive promoter, or non-promoter. (a) For complexes with 2 fragments, enrichment test captured ‘active promoter – non-promoter’ pair the most, possibly reflecting the promoter-enhancer interactions previously observed in ChIA-PET data. The third largest proportion is a contact between two active promoters. Insignificant complexes proportions are correlated with the number of non-promoters. (b) Simultaneous interactions among 3 regulatory elements can be inferred from complexes with 3 fragments. More than 50% of significant complexes involve one or two active promoter with non-promoter(s). (c) Similarly, complexes with 4 fragments are centered around active promoters, with ‘ANNN’, ‘AANN’, ‘AAAN’, and ‘AINN’ collectively making up more than 60% of the significant complexes. (d) Significant complexes had promoter fragments with higher corresponding gene expression (median=57) than insignificant complexes (median=34) (Mann-Whitney U test p-val< 2.2 × 10^-16^).

## 1 Supplementary Note

Supplementary Note 1. Extending MIA-Sig to GAM and SPRITE data

This note discusses the technical principles of the current three experimental protocols (GAM, SPRITE, and ChIA-Drop) for capturing multiplex interaction data, and the applicability of MIA-Sig to all multiplex data generated by different methods including GAM and SPRITE.

### Supplementary Note 1. Extending MIA-Sig to GAM and SPRITE data

We have developed MIA-Sig on ChIA-Drop data, but it could also potentially be suitable for denoising multiplex chromatin interactions from other methods, such as SPRITE and GAM, considering the properties of these protocols and data sets.

SPRITE uses 3-5 rounds of split-and-pool approach to barcode each chromatin complex by combinatorial indexing, with a theoretical assumption that many rounds of splitting and pooling should result in one unique barcode combination per chromatin complex. However, in practice, the split-and-pool process is limited to 4-5 rounds with a limited set of distinct barcodes, and in each round, potentially hundreds of thousands of chromatin complexes are assigned the same DNA oligo barcode. As a result, there is a certain non-zero probability of multiple complexes receiving an identical barcode combination. These unrelated complexes would be considered technical noise of SPRITE rather than specific chromatin interactions. The SPRITE technical noise profile is somewhat similar to that of ChIA-Drop of unrelated complexes partitioned in the same microfluidics droplet. Thus, MIA-sig could potentially be applied to the SPRITE data for noise assessment and data refinement.

Unlike SPRITE and ChIA-Drop, GAM captures multi-way interactions by slicing a cryopreserved nucleus and then taking one thin section to represent each nucleus for amplification and sequencing of the chromatin DNA fragments within that slice. Aggregating the sequencing data derived from slices of many nuclei, one can then identify statistically confident interactions by counting the frequency of occurrence. However, many unrelated DNA fragments could be randomly co-captured by the process, and the potential technical noise could be high. Thus, a hybrid of MIA-Sig’s distance test and the binomial test for inter-TAD contacts may be extended to GAM complexes by treating a binned fragment of a complex as a TAD.

In sum, all multiplex chromatin interaction data could have significant level of noise, and the principle nature of the noises are conceptually similar. The algorithm used in MIA-Sig considers general issues that should be applicable to all multiplex data. Although the current version of MIA-Sig is specifically developed based on ChIA-Drop data, it could be adapted to accommodate other data types. MIA-sig could thus be modified to call statistically significant complexes from SPRITE or GAM data. In addition, MIA-sig’s algorithm for calling TADs from multiplex chromatin contacts should be directly applicable to any multiplex chromatin interaction data, whether they be from ChIA-Drop, SPRITE, or GAM.

